# GATA3 mutation disrupts a functional network governed by estrogen receptor, FOXA1 and GATA3

**DOI:** 10.1101/654871

**Authors:** Motoki Takaku, Sara A. Grimm, Bony De Kumar, Brian D. Bennett, Paul A. Wade

## Abstract

Estrogen receptors (ER) are part of the nuclear receptor superfamily of transcription factors and are activated by the steroid hormone 17β-estradiol. ER forms a regulatory network in conjunction with other transcription factors, such as FOXA1 and GATA3. GATA3 has been identified as one of the most frequently mutated genes in breast cancer and is capable of specifying chromatin localization of FOXA1 and ER. How GATA3 mutations impact this transcriptional network is unknown. Here we investigate the function of one of the recurrent patient-derived GATA3 mutations (R330fs) on this regulatory network. Genomic analysis indicates that the R330fs mutant can disrupt the cooperative action of ER, FOXA1, and GATA3, and induce a change in chromatin localization of these factors. Relocalization of ER and FOXA1 is associated with altered chromatin architecture, which leads to differential gene expression in GATA3 mutant cells. These results suggest an active role for GATA3 mutants in ER positive breast tumors.

## Introduction

Cooperative action of transcription factors creates complex gene regulatory networks to maintain cell characteristics. Disruption of these regulatory networks is often associated with human diseases including cancer. Estrogen receptor alpha (hereafter ER), FOXA1 and GATA3 are essential transcription factors for mammary development, and in breast cancers, the expression levels of these three genes are highly correlated (1–7). All three are frequently expressed in luminal subtypes of breast tumors. The recent large-scale genomic profiling of breast tumors identified frequent mutations in GATA3 and FOXA1 (8–11). More than 10% of breast tumors carry GATA3 mutations, and these mutations are frequently observed in invasive ductal carcinoma. In contrast, FOXA1 mutations are more frequently detected (~7%) in invasive lobular carcinoma. In both cases, cells were found to be heterozygous for most of the mutations, with many of them located within sequences coding for their DNA-binding domains. In the case of GATA3, more than 70% of the cases are small nucleotide deletions or insertions (indel mutations), while less than 30% are missense mutations. These indel mutations induce frame-shifts, leading to protein truncation or extension. However, the functional consequences of these mutations are still largely unknown.

GATA3 and FOXA1 are known to act as pioneer transcription factors in the context of breast cancer (12–15). These pioneer factors are capable of binding closed chromatin and induce chromatin opening. Chromatin opening by pioneer factors leads to recruitment of other chromatin binding proteins (including transcription factors, remodeling factors, and histone modifiers) followed by gene activation (12,15–18). The mechanisms of these reactions remain elusive. In luminal breast cancer, GATA3 forms a transcriptional regulatory network with FOXA1 and ER, which is critical to maintain epithelial cell identity (14,19,20). Importantly, the chromatin binding activities of GATA3 and FOXA1 do not require estrogen stimulation, and, therefore, GATA3 and FOXA1 can localize at epithelial marker genes and estrogen signaling genes before ER binding (13,19). It has also been shown that chromatin localization of FOXA1 and ER is influenced by GATA3 silencing (13). However, the functional importance of GATA3 mutations for this regulatory network is unknown.

The distribution of cancer-related GATA3 mutations is focused in the C-terminal region of GATA3 (8,9,11). We previously classified those mutations, based on the location and predictive protein products, and revealed that the mutations found in the second zinc-finger domain (ZnFn2) are associated with worse patient outcomes (21). We also experimentally demonstrated that one of these mutations (R330fs: a heterozygous frame-shift mutation at arginine 330) induces alterations of GATA3 chromatin binding activities leading to more aggressive tumor phenotypes *in vitro* and *in vivo* (21). Other groups also observed similar aggressive cell characteristics in breast cancer cells that express another ZnFn2 mutation (D336fs: a heterozygous frame-shift mutation at aspartic acid 336) (22–24). The phenotypic alterations are partially explained by the action of the mutant protein (9,21,25). But the mechanism by which these changes, induced by either a two-nucleotide deletion (R330fs) or a one-nucleotide insertion (D336fs) from a single allele of GATA3 gene, become manifest remains elusive. Here, we investigate the impacts of the R330fs GATA3 mutation on the action of GATA3 co-factors, ER and FOXA1. Our genomics analysis indicates that the GATA3 mutation interrupts the regulatory network of these transcription factors by inducing redistribution of FOXA1 and ER.

## Results

### GATA3 mutation induces redistribution of ER

We previously established a GATA3 mutant cell line (CR3) by using the CRISPR-Cas9 genome editing technique (21). This cell line expresses one of the recurrent ZnFn2 mutations (R330fs) found in multiple breast cancer cohorts (8,11,26,27). The second zinc-finger domain is essential for the recognition of the GATA3 consensus motif (WGATAR). A two-nucleotide deletion from one allele induces a frame-shift and premature protein truncation (Figure 1A). Most of R330fs mutations (9 out of 10 cases) are found to be heterozygous. Therefore, this cell line mimics the situation of R330fs in human breast tumors. To investigate the impacts of the mutation on the GATA3 co-factors, we first looked at the protein expression levels of ER and FOXA1 in wild-type and mutant T47D cells. While the ER mRNA level was decreased in CR3 cells, the protein level was similar to that in wild-type T47D cells (Figure 1A). The FOXA1 expression level in the mutant cells was equivalent to wild-type cells (Figure 1A). To investigate the impacts of the GATA3 mutation on chromatin localization of these co-factors, we conducted chromatin immunoprecipitation coupled with DNA sequencing (ChIP-seq), and defined the binding peaks by HOMER (28). The peak number for ER was decreased in CR3 cells (16,872 peaks in T47D cells; 11,337 peaks in CR3 cells). We then used EdgeR (29) to define differential ER chromatin enrichment (FDR < 0.05 and |fold change| > 1.5) (Figure 1B). The majority of ER peaks (11,582 peaks, ~62%) was unchanged in CR3 cells, while ER ChIP-seq signals at 3,698 (~20%) peaks were increased, and 3,512 (~19%) ER peaks were significantly decreased (Figure 1C).

**Figure 1.**
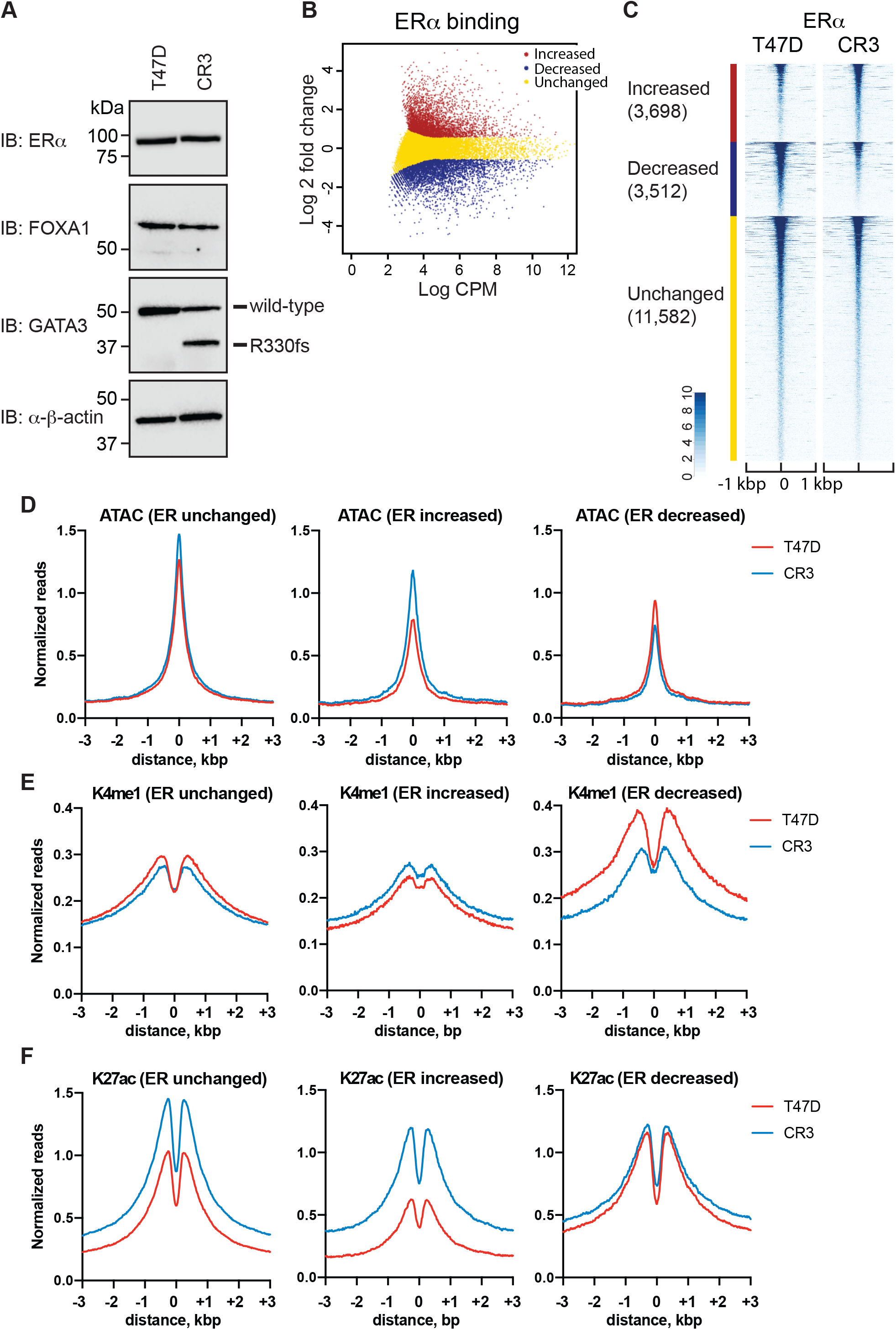
R330fs mutation induces global redistribution of ER. **(A)** Western blot showing ER, FOXA1, and GATA3 expression in T47D and CR3 cells. (**B)** Scatter plot showing differential binding of ER at ER peaks between wild-type and mutant T47D cells. CPM indicates counts per million in each peak. Increased, decreased, unchanged binding events are shown in red, blue, and yellow, respectively. (**C)** Heatmap analysis showing read density of ER ChIP-seq at ER peaks in T47D (left) or CR3 (right) cells. (**D-F)** Metaplot profiles of normalized ATAC-seq (D), H3K4me1 ChIP-seq (E), and H3K27ac (F) signals at ER ChIP-seq peaks in T47D (red) or CR3 (blue) cells.

To measure the impacts of differential binding of ER on chromatin architectures, we investigated chromatin accessibility by ATAC-seq and enhancer histone marks (H3Kme1 and H3K27ac) by ChIP-seq. Metaplot analysis of the ATAC-seq data displayed differential impacts in each peak group (Figure 1D). ER-increased binding sites showed higher ATAC-seq signals in CR3 cells, while decreased and unchanged ER peaks showed moderate reduction of ATAC-seq signals in CR3 cells. Histone H3K4me1 levels in the GATA3 mutant cells were significantly reduced at ER-decreased sites, whereas increased or unchanged peak groups displayed either slight increases or reduction in CR3 cells as compared to wild-type T47D cells (Figure 1E). Histone H3K27ac levels in CR3 cells were significantly higher at both ER-increased and unchanged peaks, while ER-decreased peaks indicated similar levels in the wild-type and mutant cells (Figure 1F). Taken together, these data suggest that the GATA3 mutation reshapes ER localization at a subset of ER binding sites leading to chromatin reprogramming.

### GATA3 mutation redirects FOXA1 chromatin localization

We next analyzed the effects of the GATA3 frameshift on FOXA1 activity. FOXA1 ChIP-seq data indicated a similar number of FOXA1 peaks in T47D (39,041 peaks) and CR3 cells (37,877 peaks). Differential peak analysis identified 4,765 increased peaks (~9%), 5,441 decreased peaks (~11%), and 40,338 unchanged peaks (~80%). Chromatin architectures were also changed at FOXA1 binding sites (Figure 2A-B). The ATAC-seq data showed a positive correlation between changes in FOXA1 binding and chromatin accessibility, consistent with its function as a pioneer transcription factor (Figure 2C). Similarly, both H3K4me1 and H3K27ac levels also correlated with differential FOXA1 binding (Figure 2D-E). FOXA1 increased binding sites showed increased levels of these active enhancer marks, while FOXA1 decreased peak regions showed decreased levels of these histone modifications. These data indicate the R330fs mutant induces redistribution of FOXA1.

**Figure 2.**
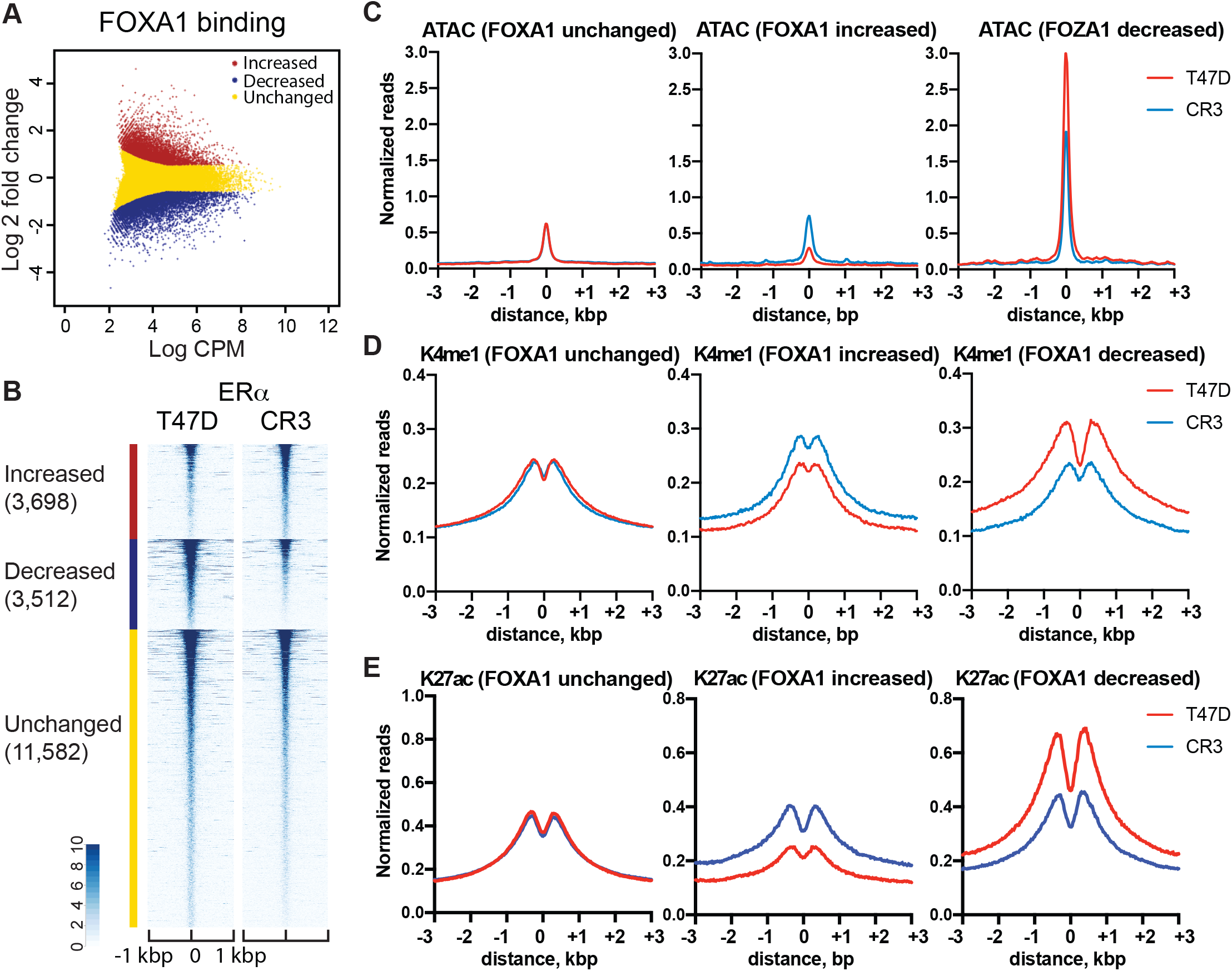
R330fs mutation reshapes FOXA1 distribution and chromatin architectures. **(A)** Scatter plot showing differential binding of FOXA1 at FOXA1 peaks in wild-type versus mutant T47D cells. CPM indicates counts per million in each peak. Increased, decreased, unchanged binding events are shown in red, blue, and yellow, respectively. (**B)** Heatmap analysis displaying normalized read density of FOXA1 ChIP-seq at FOXA1 peaks in T47D (left) or CR3 (right) cells. (**C-E)** Metaplot profiles of normalized ATAC-seq (C), H3K4me1 ChIP-seq (D), and H3K27ac (E) signals at FOXA1 ChIP-seq peaks in T47D (red) or CR3 (blue) cells.

### Redistribution of ER and FOXA1 reshapes transcriptional program

To measure the impacts of redistribution of ER and FOXA1 on gene expression, gene set enrichment analysis (GSEA) was conducted on data from cells bearing the GATA3 frame-shift mutant. We first assigned the ER and FOXA1 peaks to nearest genes (within 50 kbp). The genes close to sites of ER-increased peaks in CR3 cells were frequently up-regulated when compared to wild-type T47D cells, whereas ER-decreased peaks are often observed near down-regulated genes (Figure 3A). Similar to the situation with ER, differential binding of FOXA1 exhibited strong positive correlation with gene expression changes in wild-type versus mutant T47D cells (Figure 3B). The gene ontology analysis of ER-or FOXA1-associated genes indicated that sites containing either ER or FOXA1 decreases in peaks were enriched for pathways related to gland development, epithelial cell development, and hormone signaling (Figure 3C, Supplementary figure 2A). Increased binding of ER was associated with cell adhesion pathways, while FOXA1-increased sites were enriched in genes related to reproductive system development (Figure 3F, Supplementary figure 2A). To dissect the upstream regulators of differentially expressed genes that are associated with ER or FOXA1 differential binding, IPA (Ingenuity Pathway Analysis, QIAGEN) pathway analysis was conducted. This upstream regulator analysis indicated that loss of ER and FOXA1 peaks were associated with down-regulation of signaling cascades regulated by ligand-dependent nuclear receptors such as PPARG (Figure 3E, Supplementary figure 2C). TGFB1 pathway-related genes were also enriched in both cases. On the other hand, increased binding of ER was preferentially found at genes involved in AHR and CCL5, and these pathways were predicted to be activated (Figure 3F). On the other hand, increased FOXA1 peaks were significantly associated with TP53, CD24, and TNF pathway activation (Supplementary figure 2D). Taken together, these results suggest that relocalization of ER and FOXA1 by the GATA3 R330fs mutant contribute to cellular reprogramming at the transcription level and to more aggressive phenotypic alterations in CR3 cells.

**Figure 3.**
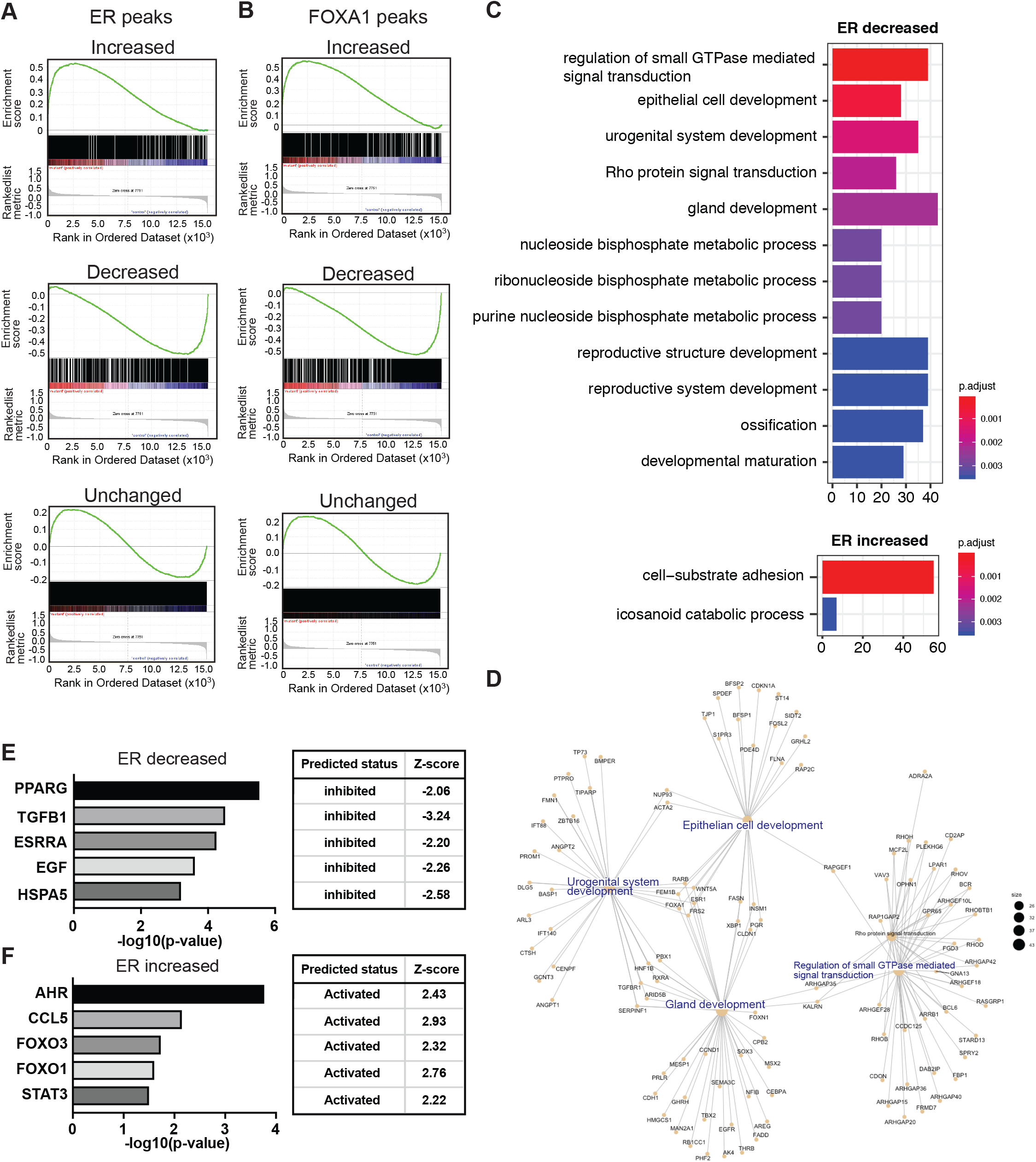
Relocalizations of ER and FOXA1 are associated with differential gene expression. **(A)** GSEA analysis indicating correlation between ER binding and gene expression in each ER peak group. **(B)** GSEA analysis indicating correlation between FOXA1 binding and gene expression in each FOXA1 peak group. **(C)** Biological pathway analysis of genes associated with ER decreased (top) or increased (bottom) peaks. Top 12 significantly enriched pathways are shown. **(D)** Gene networks enriched at ER decreased peaks. Functional enrichment analysis was conducted using genes associated with ER-decreased peaks. **(E)** Upstream regulator analysis of genes associated with ER decreased binding. **(F)** Upstream regulator analysis of genes associated with ER increased peaks.

### GATA3 mutant disrupts ER-FOXA1-GATA3 network

To further explore the function of the GATA3 frame-shift mutant, we next focused on cooperative action between ER, FOXA1, and GATA3 on chromatin. Consistent with previous studies, these three transcription factors are colocalized at a subset of binding sites (Figure 4A-B) (13,19). In the GATA3 mutant cells, peak overlap frequency of ER, FOXA1, and GATA3 was slightly decreased, but the colocalization was still detected. To characterize the cooperative binding sites, we classified GATA3 peaks identified in the wild-type T47D cells into 4 groups based on the ChIP-seq signals of ER, FOXA1, and GATA3 as follows: EFG: ER^+^, FOXA1^+^, GATA3^+^; EG: ER^+^, FOXA1^−^, GATA3^+^; FG: ER^−^, FOXA1^+^, GATA3^+^; G alone: ER^−^, FOXA1^−^, GATA3^+^.. Interestingly, chromatin features were clearly different in each binding group. EFG sites possessed higher ATAC-signals as well as more enhancer marks (H3K4me1 and H3K27ac) as compared to the other peak groups (Figure 4C). The genes associated with EGF peaks were significantly enriched for cell junction, mammary gland development, and reproductive system development, suggesting that the ER-FOXA1-GATA3 cooperative binding is essential to maintain luminal cell identity (Supplementary figure 4). In the GATA3 frame-shift mutant cells, binding of wild-type GATA3 was significantly decreased at EGF sites, while GATA3 alone peaks exhibited increased binding of wild-type GATA3 (Figure 4D, Supplementary figure 2). In addition, binding of both ER and FOXA1, chromatin accessibility, and active enhancer marks were all reduced at EGF sites (Figure 4B, 4E-F).

**Figure 4.**
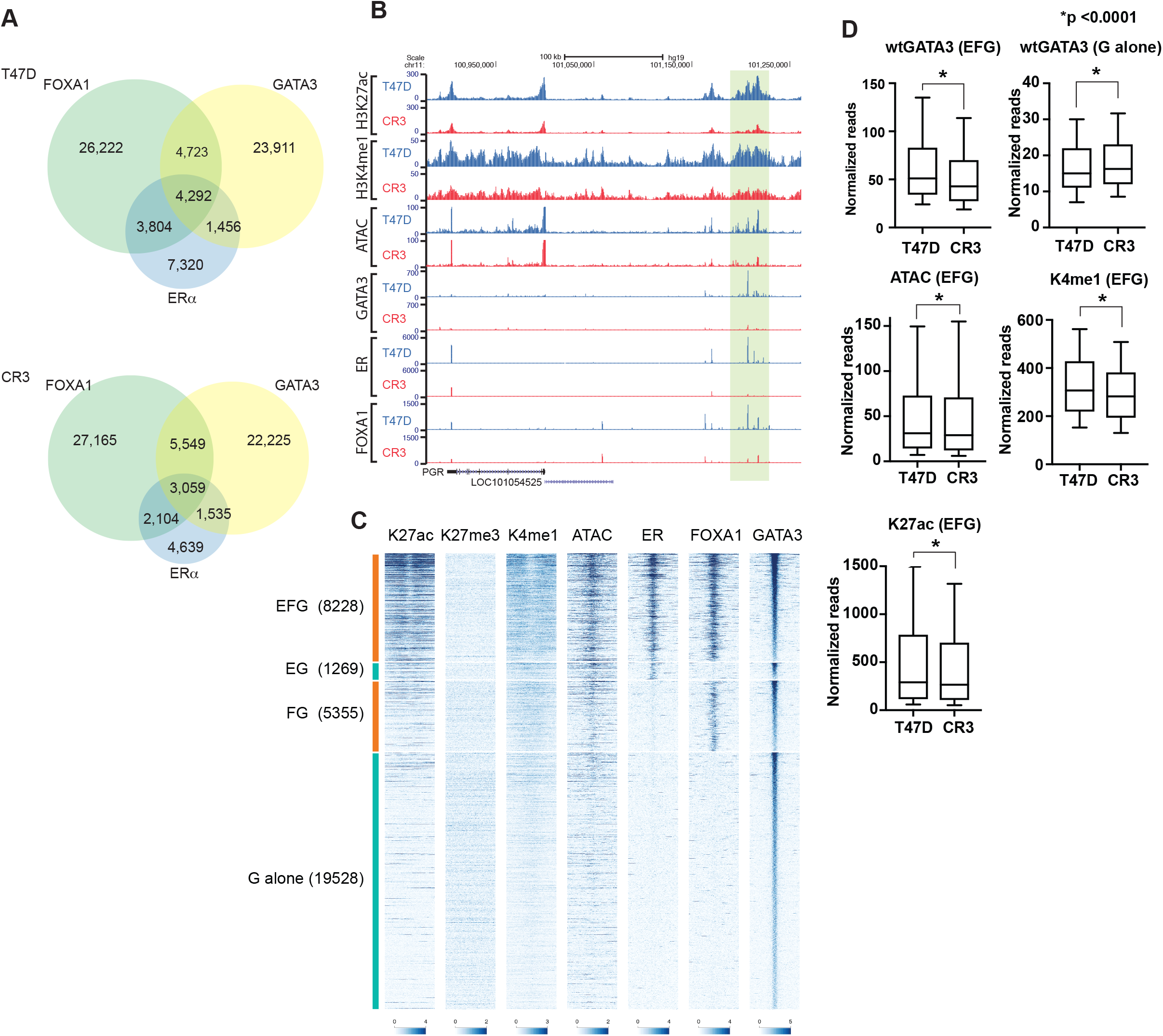
R330fs mutant interrupts ER-FOXA1-GATA3 network. **(A)** Venn diagram showing overlap between ER, FOXA1, and GATA3 peaks in T47D cells (top) and CR3 cells (bottom). **(B)** Representative genome browser tracks at ER-FOXA1-GATA3 co-localized locus. **(C)** Heatmap showing H3K27ac, H3K27med, H3K4me1, ATAC, ER, FOXA1, and GATA3 ChIP-seq in each GATA3 peak category. **(D)** Box plot showing normalized read of GATA3 ChIP-seq at EFG or_G peaks in T47D (left) or CR3 (right) cells. Y-axis indicates normalized reads per peak. **(E)** Box plot showing normalized read of ER and FOXA1 ChIP-seq at EFG or_G peaks. **(F)** Box plot showing normalized read of ATAC-seq, H3K4me1, and H3K27ac ChIP-seq signals.

To confirm the reduction of cooperative binding, we also looked at the distribution of wild-type GATA3 at differential ER and FOXA1 binding peaks (defined above). A significant reduction of GATA3 ChIP signals was detected at ER-decreased peaks in CR3 cells, while a moderate GATA3 increase was detected at ER-increased peaks in CR3 cells (Figure 5A). Similarly, GATA3 ChIP signals were also drastically reduced at FOXA1-decreased peaks in the mutant cells (Figure 5B).

**Figure 5.**
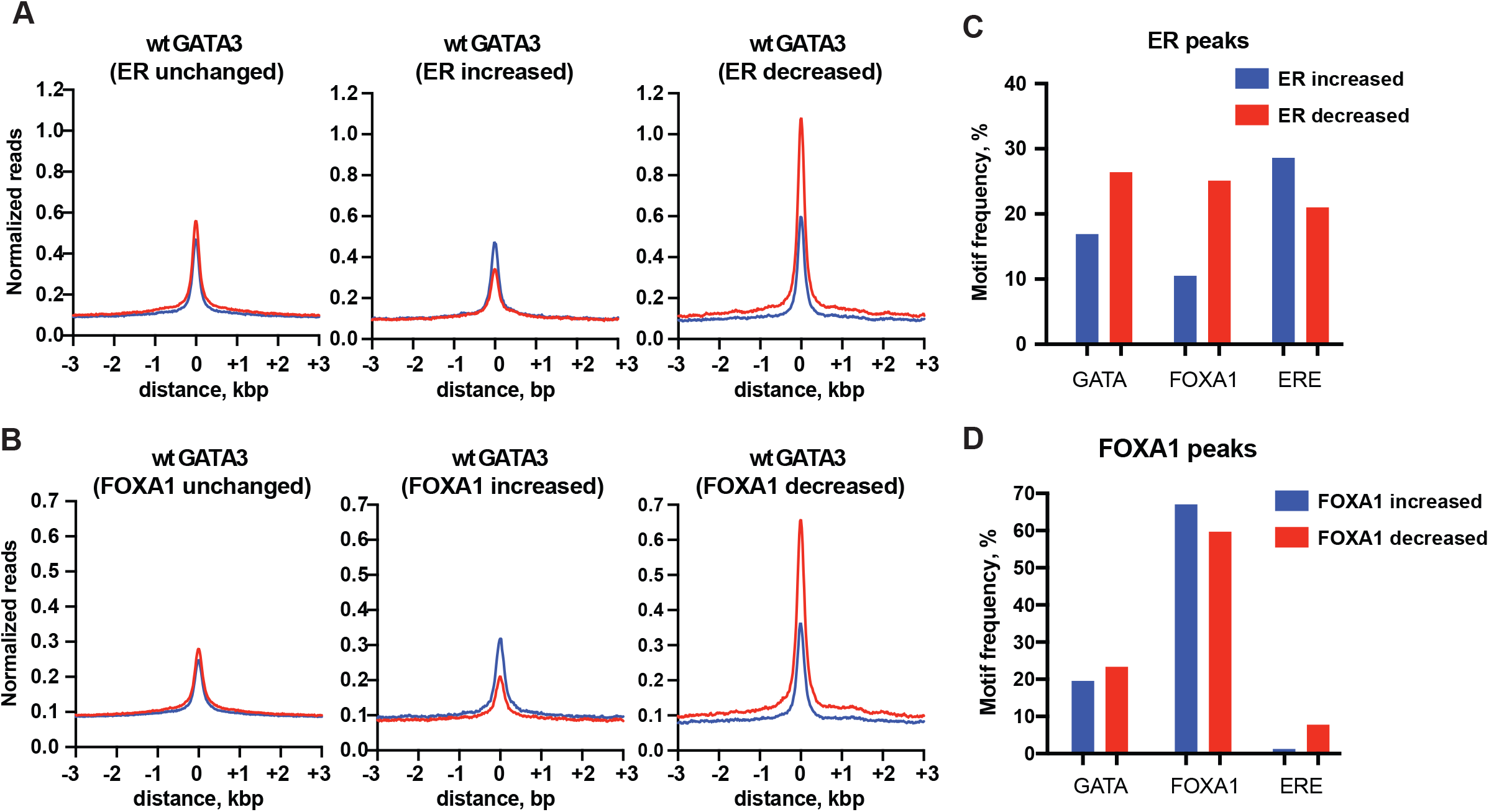
Motif analysis suggests decreased cooperation of ER-FOXA1-GATA3 complex in GATA3 mutant cells. **(A)** GATA3 binding profiles at ER differential binding peaks. GATA3 ChIP-seq data from T47D cells is shown in red, and the profile from CR3 cells is shown in blue. **(B)** GATA3 binding profiles at FOXA1 differential binding peaks. **(C)** GATA3, FOXA1, and ER motif frequency in ER increased or decreased binding sites. **(D)** GATA3, FOXA1, and ER motif frequency in FOXA1 increased or decreased binding sites.

To gain further insights into the mechanisms underlying the GATA3 mutant-induced relocalization of FOXA1 and ER, we analyzed the motif frequency at differential binding sites. While the metaplot analysis displayed a weak increase of GATA3 ChIP-seq signals at peaks showing an increase in FOXA1 or ER, the GATA3 consensus motif (WGATAR) was less frequently detected at these peaks as compared to ER and FOXA1 increased peaks (Figure 5C). Similarly, peaks associated with increases in FOXA1 binding in CR3 cells carry less ER motifs than peaks associated with less FOXA1, and ER increased peaks manifest less FOXA1 motifs than do ER decreased peaks. However, their own motifs are more frequently observed in increased peaks than decreased peaks. Collectively, these data suggest that the GATA3 mutant disrupts the gene regulatory hub governed by ER, FOXA1 and GATA3.

## Discussion

Somatic mutations within transcription factors are one of the common phenomena in cancer. One such example, GATA3, has been identified as one of the most frequently mutated genes in breast cancer. The biological significance of GATA3 mutations remains unclear. We previously established a GATA3 mutant cell line that endogenously expresses one of the recurrent GATA3 heterozygous mutants (R330fs) (21). This cell line exhibits more aggressive phenotypes at both cellular and transcriptome levels. The R330fs mutation alters distribution of wild-type GATA3 on chromatin leading to down-regulation of the progesterone receptor signaling pathway. R330fs mutant also shows a unique chromatin binding pattern as compared to wild-type. However, we observed more than a thousand differentially expressed genes between wild-type and mutant GATA3 cells, suggesting other potential mechanisms that may contribute to the drastic phenotypic changes observed in GATA3 mutant cells. Our present study demonstrates that the R330fs mutant is sufficient to interrupt and reshape chromatin localization of two GATA3 co-cofactors, ER and FOXA1. Redistribution of ER and FOXA1 is associated with changes in chromatin architectures and gene expression. Decreased binding of ER, FOXA1, and GATA3 was predominantly observed within those loci where three factors are co-localized. These co-localizations are enriched at the sites close to epithelial marker genes including KRT18, CLND1, and OCLN. The R330fs mutant cells exhibit partial epithelial-to-mesenchymal transition (21). Therefore, impaired binding at the co-occupied sites may cause reduction of epithelial marker genes in CR3 cells (ref).

The mechanism of co-factor relocalization in mutant cells needs further investigation. It has been shown that silencing of GATA3 changes localization of ER and FOXA1, and of enhancer activities that interact with ER-dependent genes (13). The R330fs mutation can induce both loss and gain of ER, FOXA1, and GATA3 binding, whereas GATA3 binding at ER increased sites showed moderate induction. Consistently, a moderate increasement of ER binding was observed at GATA3 increased sites, suggesting that the functional correlation between GATA3 and ER is damaged by the R330fs mutant. Consistently, the frequency of the GATA3 consensus motifs are clearly decreased at ER increased sites as compared to ER decreased sites. In addition, the frequency of the peaks that contain both ER and FOXA1 motifs are also significantly decreased at ER and FOXA1 increased sites. Decreased number of GATA3 motifs at ER or FOXA1 increased binding sites also implies a tethering mechanism of GATA3 by ER or FOXA1 (30). Together, these data suggest that the R330fs mutation compromises the regulatory network governed by ER, FOXA1, and GATA3.

Importantly, sites jointly bound by ER, FOXA1 and GATA3 showed strong open chromatin features as well as active enhancer status. Our GATA3 mutant cell line model suggests that intact binding of the three factors are essential to maintain a highly active chromatin state at these loci. Therefore, these chromatin regions might be under dynamic competition between ER-FOXA1-GATA3 gene activators and other chromatin repressors (31–33), and the altered DNA binding ability of the R330fs mutant weakens the overall chromatin binding affinity of the ER-FOXA1-GATA3 complex (21,25). Although the GATA3 mutations found in breast cancers are tend to accumulate at the C-terminal region, the protein products produced by these mutations (e.g. frame-shift mutations and missense mutations) are not identical and likely have differential impacts on the GATA3 protein activity. Defining molecular functions of other mutants would be useful to further understand the biological significance of breast cancers carrying the mutations in this gene.

## Materials and methods

### Cell line and cell culture

The T47D cell line was originally purchased from ATCC. The CR3 GATA3 mutant cell clone (T47D ^R330fs/wild-type^) was previously established by using CRISPR-Cas9 based gene editing technique (21). T47D cells that express FLP recombinase was used as GATA3 wild-type T47D control. All cells were grown in DMEM high-glucose medium supplemented with 10% fetal bovine serum (Thermo Fisher Scientific) at 37°C with 10% CO_2_.

### ChIP-seq

The details of the procedures were previously described (12,21). After removing medium, cells were fixed with formaldehyde (4% in PBS) at room temperature for 10 min, and stored at −80 °C. Fixed cells were thawed on ice and treated with hypotonic buffer containing 10 mM HEPES-NaOH pH 7.9, 10 mM KCl, 1.5 mM MgCl2, 340 mM sucrose, 10% glycerol, 0.5% Triton X-100, and Halt Protease & Phosphatase Single-Use Inhibitor Cocktail (Thermo Fisher Scientific) for 5 min. The cells were further resuspended in lysis buffer containing 20 mM Tris-HCl pH 8.0, 2 mM EDTA, 0.5 mM EGTA, 0.5 mM PMSF, 5 mM sodium butyrate, 0.1% SDS, and protease inhibitor cocktail. Chromatin was sonicated using a Covaris S220. Immunoprecipitation was performed with anti-ER-alpha (Santa Cruz, HC-20), anti-FOXA1 (abcam, ab5089), anti-H3K4me1 (abcam ab8895), anti-H3K27ac (abcam ab4729), or anti-H3K27me3 (abcam ab192985) antibody. The sequencing libraries were prepared by the NEXTflex Rapid DNA-seq kit (Bioo Scientific Corporation). All sequencing was conducted at the NIEHS Epigenomics Core Facility (NextSeq 500 or NovaSeq 6000 NGS system, Illumina). Reads were filtered based on a mean base quality score >20, and mapped to hg19 genome by Bowtie 0.12.8 (34), and non-duplicate reads were used for the analysis.

### Peak analysis

ChIP-seq peaks were identified by the HOMER software using the default parameters (28). Differential ER or FOXA1 binding events between GATA3 wild-type and mutant T47D cells were identified by EdgeR with FDR < 0.05 and |fold change| > 1.5 as described in (21). Read counts within a 400 bp window centered on each peak were collected from ChIP-seq data in wild-type T47D cells or CR3 cells. For the motif frequency calculation, the HOMER tool perl script (findMotifs.pl) was used to find each motif within the peaks (400 bp window). FOXA1(Forkhead)/MCF7-FOXA1-ChIP-Seq(GSE26831)/Homer (Motif 88), ERE(NR),IR3/MCF7-ERa-ChIP-Seq(Unpublished)/Homer (Motif 71), or WGATAR was used to detect each motif.

### Transcriptome analysis

For the GSEA analysis, ChIP-seq peaks were associated with closest genes within 50 kbp from the nearest TSS. The same gene lists were used to conduct IPA pathway analysis. Functional analysis of ER or FOXA1 peak associated genes was performed in Bioconductor (version 3.8) using package clusterProfiler (version 3.5) for the enrichment of ontology related to the biological process (35). P-value was adjusted using the BH (Benjamini-Hochberg) method. Enriched pathways with an adjusted p-value less than 0.01 and q-value less than 0.05 were reported. Top 12 Enriched ontology terms are shown as Bar plot.

### Data accession

The RNA-seq, ChIP-seq, and ATAC-seq data have been deposited in NCBI’s Gene Expression Omnibus and are accessible under accession number GSE99479 (https://www.ncbi.nlm.nih.gov/geo/query/acc.cgi?acc=GSE99479).

## Acknowledgement

We gratefully acknowledge NIEHS/NIH core groups (Epigenomics core and Integrated Bioinformatics core) for outstanding technical assistance. We thank Dr. John D. Roberts for insightful suggestions on the manuscript. This work was supported, in part, by the Intramural Research Program of the National Institute of Environmental Health Sciences, NIH (ES101965 to PAW), Division of Intramural Research Innovative Research Award (DIRA) from NIEHS (to MT), and by the start-up fund provided by the University of North Dakota School of Medicine and Health Sciences, Department of Biomedical Sciences (to MT).

---

**Supplementary figure S1.**
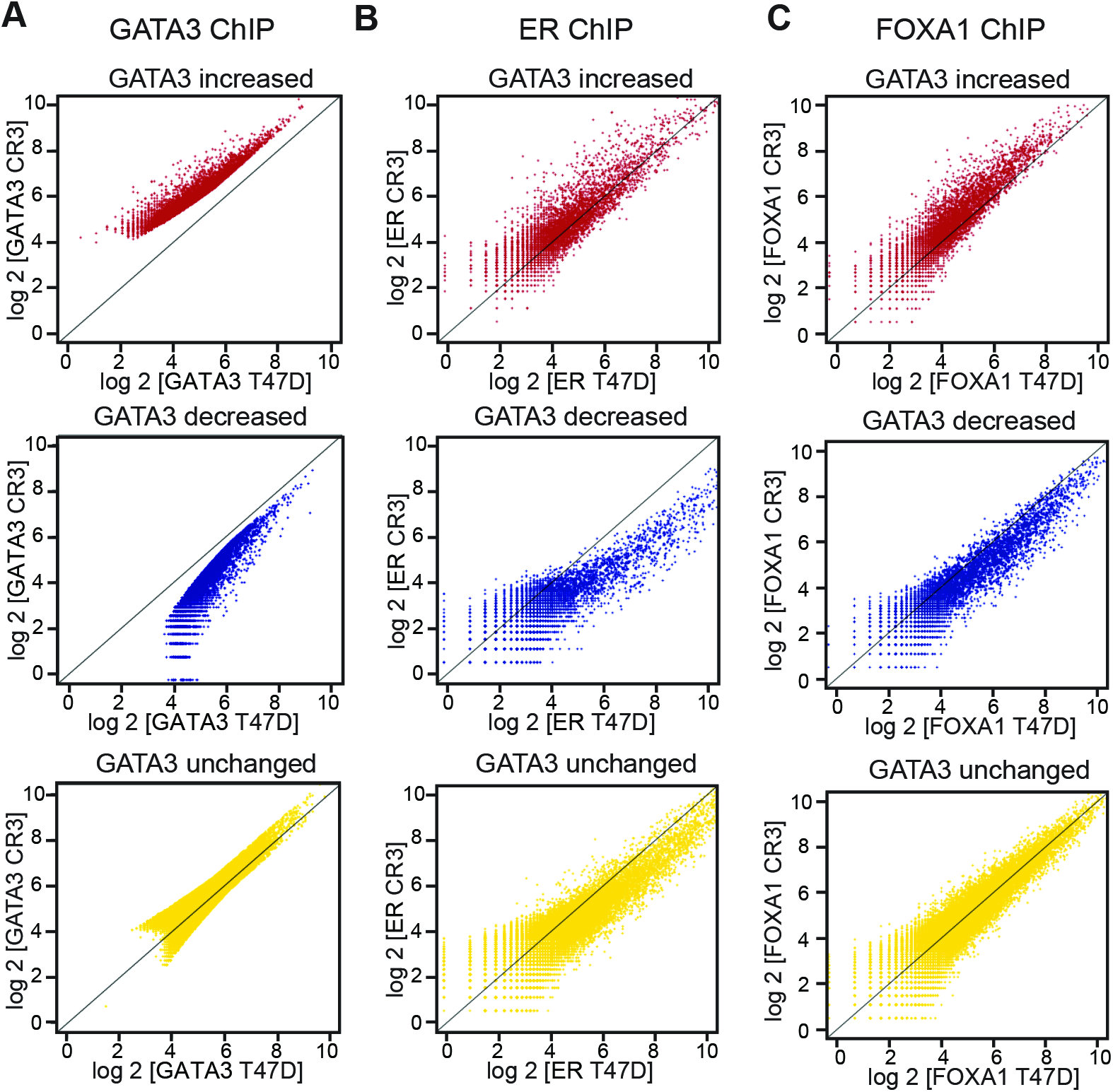
Differential binding of GATA3 in CR3 cells is correlated with ER and FOXA1 redistribution **(A)** Scatter plot showing GATA3 ChIP-seq signals in each differential GATA3 binding group. X-axis indicates GATA3 ChIP-seq in wild-type T47D cells, while Y-axis indicates GATA3 ChIP-seq in CR3 cells. **(B)** Scatter plot showing ER ChIP-seq signals in each GATA3 peak group. X-axis: ER ChIP-seq in wild-type T47D cells, Y-axis: ER ChIP-seq in CR3 cells **(C)** Scatter plot showing FOXA1 ChIP-seq signals in each GATA3 peak group. X-axis: FOXA1 ChIP-seq in wild-type T47D cells, Y-axis: FOXA1 ChIP-seq in CR3 cells.

**Supplementary figure S2.**
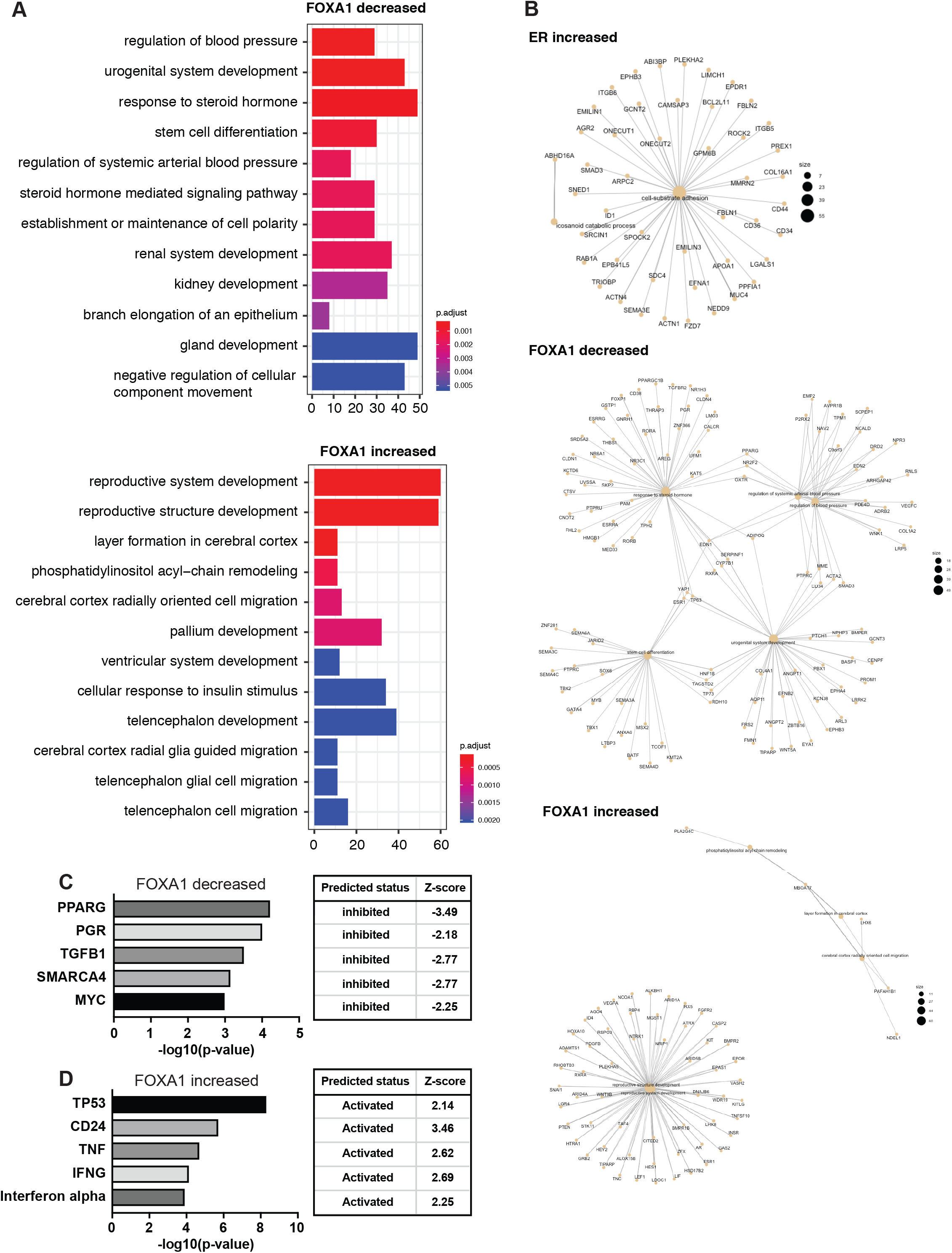
Functional enrichment analysis at differential ER and FOXA1 binding. **(A)** Barplots showing top 12 enriched biological pathways that are associated with FOXA1 decreased (top) or increased (bottom) binding sites. **(B)** Gene networks enriched at ER decreased (top), FOXA1 decreased (middle), and FOXA1 increased (bottom) peaks. **(C)** Upstream regulator analysis of genes associated with FOXA1 decreased binding. **(D)** Upstream regulator analysis of genes associated with FOXA1 increased peaks.

**Supplementary figure S3.**
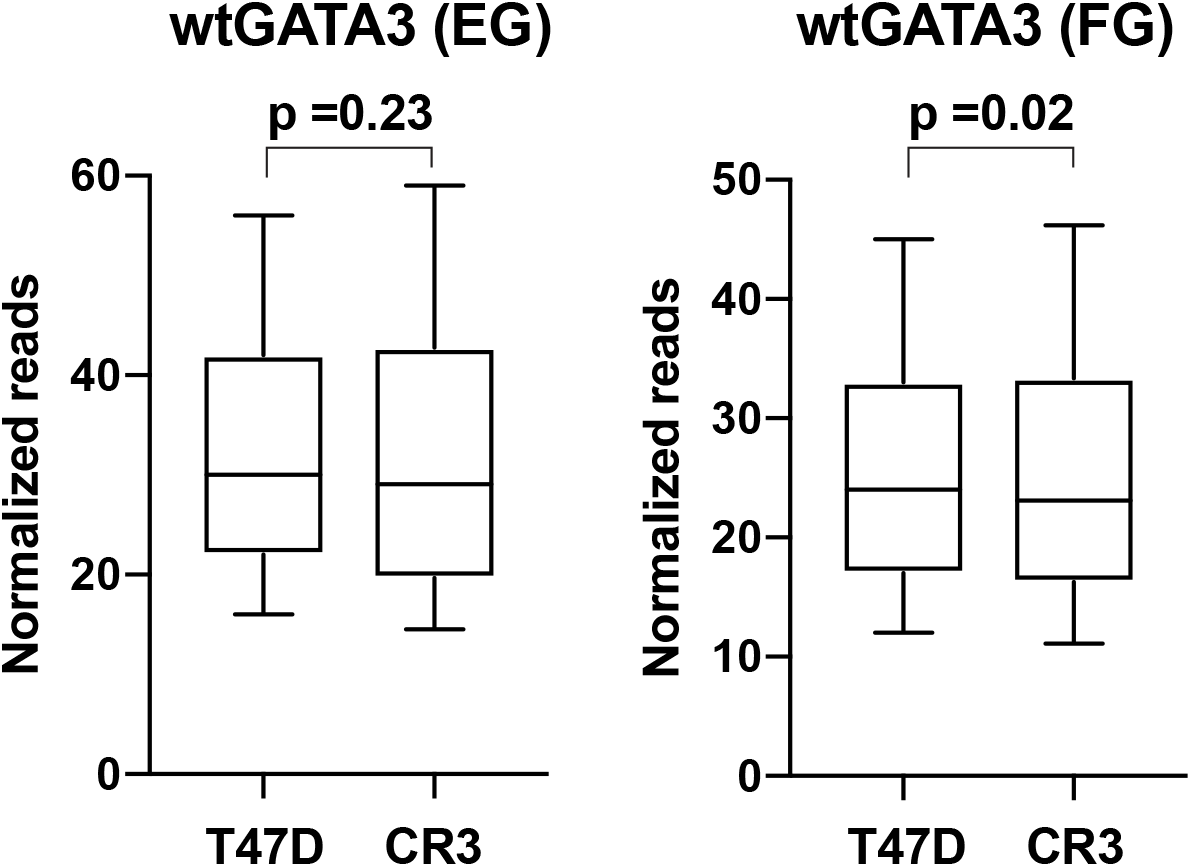
Box plot showing normalized read of GATA3 ChIP-seq at EG or FG peaks in T47D (left) or CR3 (right) cells. Y-axis indicates normalized reads per peak.

**Supplementary figure S4.**
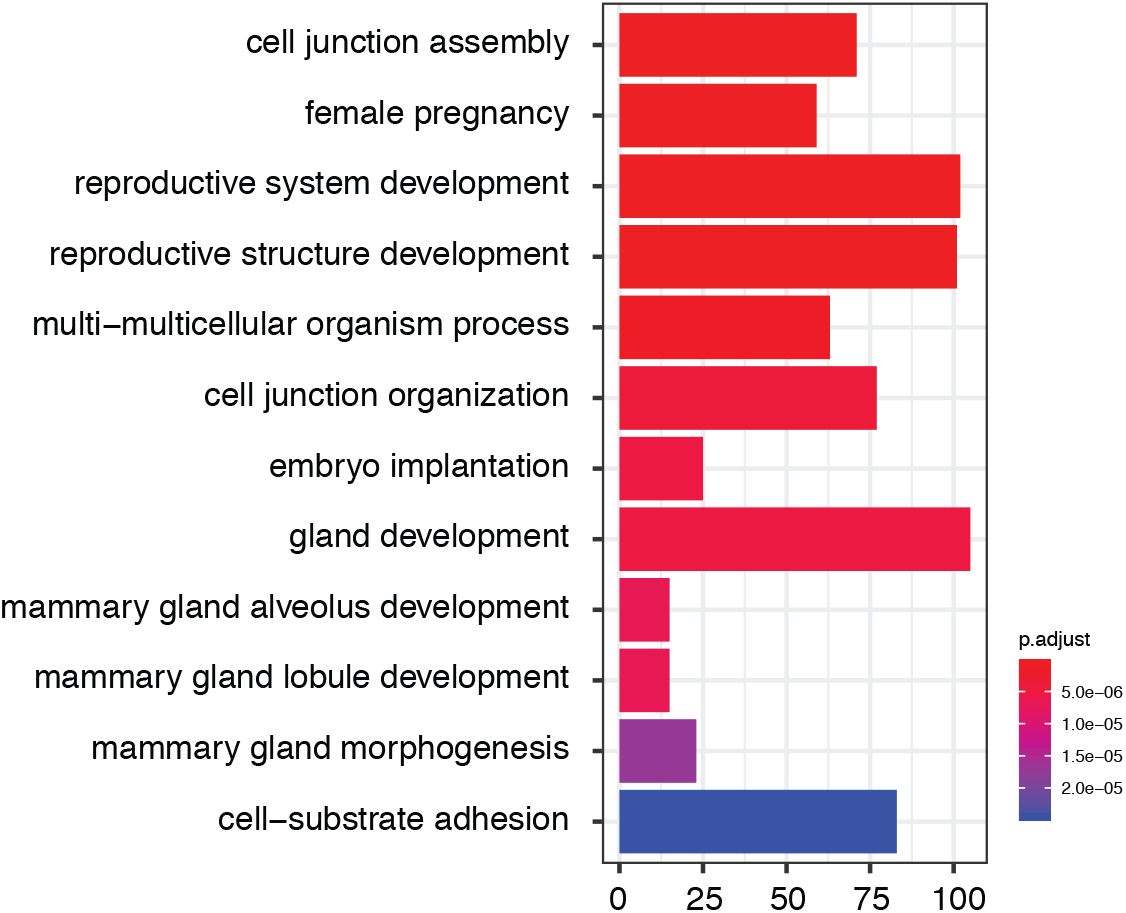
Barplot showing top 12 biological pathways that are significantly enriched at EFG peaks.

